# Semaphorin 3A promotes resilience of sleep and cognition to brain injury

**DOI:** 10.64898/2026.06.24.734382

**Authors:** D. Necula, Y. Voskobiynyk, S. Poluri, J.T. Paz

## Abstract

Chronic sleep disruption and cognitive deficits are debilitating long-term consequences of traumatic brain injury (TBI), yet the endogenous mechanisms that drive network-level recovery remain poorly understood. Here, we demonstrate that during the chronic phase post-injury, network oscillations critical for memory, including slow oscillations, delta waves, and sleep spindles, undergo precise adaptive tuning that sustains sleep-dependent memory consolidation. This functional circuit resilience requires Semaphorin-3A (Sema3a); loss of Sema3a function prevents the protective reorganization of non-rapid eye movement (NREM) sleep architecture, impairs sleep-dependent memory consolidation, and exacerbates cortical lesion size. Together, these findings identify Sema3a as an innate protective mechanism that preserves sleep architecture and cognitive function following TBI.

## INTRODUCTION

Sleep disruption is among the most common long-term consequences of traumatic brain injury (TBI), affecting 30–70% of patients^1^. Sleep disturbances can be disabling, leading to impaired cognition^2^. However, while many individuals develop chronic sleep impairments, others show only mild or transient symptoms, suggesting the presence of endogenous protective mechanisms^2^. Identifying such mechanisms may point to new therapeutic strategies for a condition that lacks effective treatments.

Semaphorins may serve as one such candidate. This class of proteins is best known for its role in axon guidance during development, but also regulates adult neuroplasticity in injury and disease^3^. In particular, Semaphorin-3A (Sema3a), a secreted guidance cue that signals through a receptor complex composed of neuropilin-1 (Nrp1), A-type plexins, and integrins, has been shown to contribute to adult neuroplasticity^4^. Namely, disruption of Sema3a–Nrp1 signaling via the Sema3a K108N point mutation prevents neurons from restoring neurotransmission following postsynaptic receptor blockade^4^. This form of neuroplasticity, for which Sema3a is required, has also been shown to oppose the progressive neurodegeneration seen in mouse models of amyotrophic lateral sclerosis^5^. Because TBI similarly imposes a profound perturbation on neuronal circuits, Sema3a-dependent plasticity may represent an endogenous mechanism promoting the resilience of sleep after brain injury.

Here, we used Sema3a ^K108N^ mutant mice, hereafter referred to as Sema3a mutant mice, to test whether Sema3a contributes to adaptive changes in sleep architecture, sleep-dependent memory consolidation, and spontaneous behavior following controlled cortical impact (CCI) injury to the right primary somatosensory cortex (S1). Our findings identify Sema3a as a critical mediator of sleep adaptations following TBI and support a role for Sema3a in promoting resilience to chronic neurological dysfunction after brain injury.

## RESULTS

### Sema3a is required for adaptive tuning of NREM architecture after TBI

Coordinated network rhythms that occur in non-rapid eye movement (NREM) sleep, including slow oscillations (SOs), delta waves, and sleep spindles, are emergent properties of highly interconnected, distributed neural circuits that underlie cognitive processes like memory consolidation^6–12^. Because these synchronized rhythms rely on widespread structural and functional connectivity, they are uniquely vulnerable to cortical injury^12–16^. Consequently, post-traumatic alterations in these global NREM rhythms may serve as a functional readout of overall network integrity and functional recovery^12–16^.

To assess the chronic effects of the TBI on NREM architecture, we performed electrocorticography (ECoG) ipsilateral and contralateral to the injury site 1.5 months post-injury (**Figure 1A and 1B).** Overall, the fraction of time spent in NREM did not differ significantly between groups (**Figure S1A and S1B**), though TBI prolonged the mean duration of NREM bouts regardless of genotype (**Figure S1C**).

**Figure 1.**
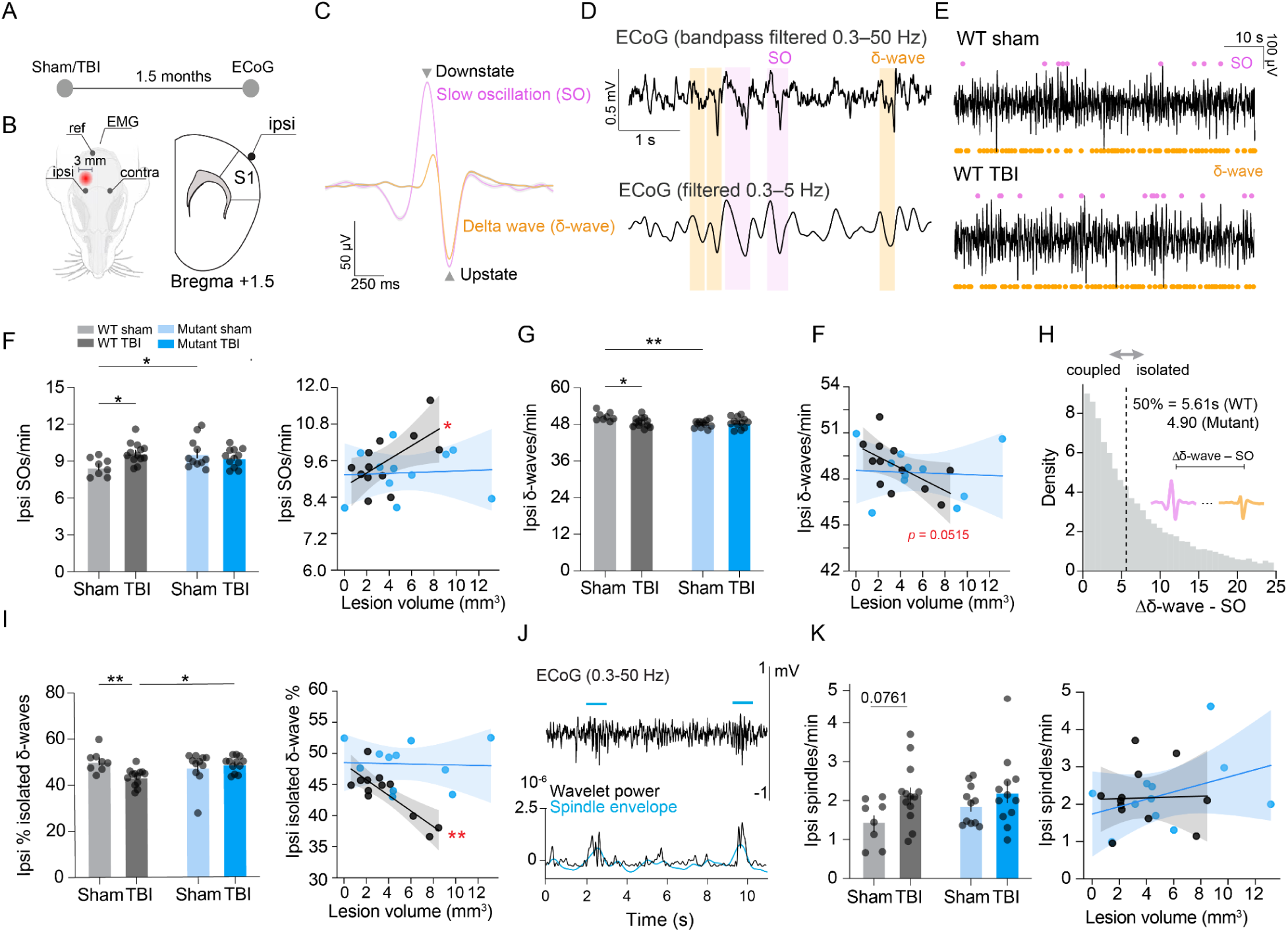
Sema3a is required for adaptive tuning of NREM architecture after TBI. **(A)** Experimental timeline. **(B)** Schematic of ECoG setup and coronal section indicating placement of the ipsilateral (ipsi) primary somatosensory cortex (S1) electrode. The contralateral (contra) electrode is symmetrical in the opposing hemisphere (not shown). The red area indicates the site of TBI. Image created with Biorender.com. **(C)** Average slow oscillation (SO) and delta wave (δ-wave) from a representative mouse. **(D)** Representative SOs and δ-waves in a 5-s NREM segment filtered at 0.3–50 Hz (top) and in the delta band at 0.3–5 Hz (bottom). **(E)** Representative NREM traces from a WT sham and a WT TBI mouse. **(F–G)** SO (F) and delta wave (G) rate in NREM on the ipsi hemisphere with lesion volume correlations. WT sham, N = 8; WT TBI, N = 14; Sema3a mutant sham, N = 11; Sema3a mutant TBI, N = 13. Two-way ANOVA with Holm–Sidak multiple comparisons correction. Simple linear regression for correlations. Asterisks next to each regression line indicate a significant *p*-value. See Table 1 for linear regression statistics. **(H)** Histogram of the temporal delay between SO and δ-wave upstates in the WT sham. Median for WT sham = 5.61 s; median for Sema3a mutant sham = 4.90 s. N’s and statistics as in F and G. **(I)** Percent of isolated or coupled ipsi δ-waves with lesion volume correlation. N’s and statistics as in F and G. **(J)** Spindle detection using the Morlet wavelet approach. ECoG filtered 0.3–50 Hz (top), Morlet wavelet power with the Hilbert transform spindle envelope superimposed (bottom). Blue bars indicate detected spindles. **(K)** Ipsi spindle rate in NREM with lesion volume correlation. WT sham, N = 8; WT TBI, N = 13; Sema3a mutant sham, N = 11. Sema3a mutant TBI, N = 12. Statistics as in F, G, and I. See Table 1 for linear regression statistics. \**p* < 0.05, \*\**p* < 0.01. Error bars show mean ± SEM across animals.

**Table 1.** Linear regression statistics for data in Figure 1F,G,I,K and Supplementary Figure 2A–D.

|  |  |  |
| --- | --- | --- |
| Contra SOs/min | WT TBI | $r^2 = 0.25, p = 0.0997$ |
| | Mutant TBI | $r^2 = 0.01, p = 0.793$ |
| Ipsi SOs/min | WT TBI | $r^2 = 0.37, *p = 0.0352$ |
| | Mutant TBI | $r^2 = 0.00, p = 0.861$ |
| Contra δ-waves/min | WT TBI | $r^2 = 0.39, *p = 0.0293$ |
| | Mutant TBI | $r^2 = 0.00, p = 0.943$ |
| Ipsi δ-waves/min | WT TBI | $r^2 = 0.39, p = 0.0515$ |
| | Mutant TBI | $r^2 = 0.00, p = 0.858$ |
| Contra isolated δ-wave % | WT TBI | $r^2 = 0.64, **p = 0.0018$ |
| | Mutant TBI | $r^2 = 0.01, p = 0.812$ |
| Ipsi isolated δ-wave % | WT TBI | $r^2 = 0.69, **p = 0.001$ |
| | Mutant TBI | $r^2 = 0.00, p = 0.904$ |
| Contra spindles/min | WT TBI | $r^2 = 0.00, p = 0.997$ |
| | Mutant TBI | $r^2 = 0.2, p = 0.172$ |
| Ipsi spindles/min | WT TBI | $r^2 = 0.00, p = 0.915$ |
| | Mutant TBI | $r^2 = 0.16, p = 0.229$ |

To assess how Sema3a and TBI might contribute to sleep architecture, we next quantified the rate of SOs, delta waves, and sleep spindles during NREM.

Within the delta frequency band in NREM, SOs and delta waves predominate^7^ (**Figure 1C and 1D**) and perform functionally antagonistic roles in memory consolidation, with SOs thought to promote it and delta waves thought to weaken it ^7,12,17–22^.

Sleep spindles are 9–15 Hz NREM oscillations crucial for memory consolidation^7,23–25^, particularly through their interaction with SOs and delta waves^7,8,10,12^. Moreover, sleep spindle activity is also associated with improved outcomes after TBI^26,27^.

If Sema3a serves a neuroprotective function in TBI, we would expect it to bias NREM towards an oscillatory state optimal for memory and cognition (namely, more SOs and spindles, and fewer delta waves).

Consistent with this hypothesis, WT TBI mice exhibited a significant increase in bilateral SO rate relative to WT sham controls, with ipsilateral SO rate correlating with lesion volume (**Figure 1C–F, and Figure S2A**). WT TBI mice also demonstrated a small yet significant bilateral reduction in delta wave rate, which correlated with lesion volume in the contralateral hemisphere (**Figure 1G and Figure S2B)**. By contrast, Sema3a mutant TBI mice had fewer contralateral SOs compared to Sema3a mutant sham mice, and no significant changes in ipsilateral SO rate or bilateral delta wave rate (**Figure 1F,1G and Figure S2A, S2B**). Furthermore, Sema3a mutant TBI mice exhibited no correlation between lesion volume and SO or delta wave rate, whether contralateral or ipsilateral (**Figure 1F,G and Figure S2A, S2B**).

Delta waves tend to occur in close temporal proximity to SOs, and their decoupling can impede memory consolidation^12^. In WT TBI mice, we observed a bilateral reduction in the proportion of isolated delta waves (defined as delta waves occurring beyond the median SO–delta wave temporal delay) that correlated with lesion volume (**Figure 1H,1I and Figure S2C)**. Thus, rather than a uniform depletion of delta waves, WT TBI mice preferentially lose SO-decoupled delta waves—likely reflecting the increased rate of SOs after injury. No change was observed in the proportion of isolated delta waves in Sema3a mutant TBI mice on either the ipsilateral or contralateral hemisphere, nor did they exhibit any correlation between isolated delta wave proportion and lesion volume (**Figure 1H,1I and Figure S2C**). Notably, Sema3a mutant sham mice consistently demonstrated baseline differences in bilateral SO and delta wave rate (**Figure 1F and 1G)**.

We also observed a trend toward higher ipsilateral sleep spindle rate and a significantly higher contralateral sleep spindle rate following TBI in WT mice, neither of which was observed in Sema3a mutant TBI mice (**Figure 1J,K and Figure S2D**). Unlike SOs, delta waves, and isolated delta waves, spindle density did not correlate with lesion volume regardless of hemisphere (**Figure 1K and Figure S2D**).

A known feature of spindles is that they tend to peak at the upstate of both SOs and delta waves, representing a period of synchronous neuronal depolarization (**Figure S3A and S3B**) ^7,12,28^. This interaction is crucial for memory: optogenetic decoupling of the SO upstate and spindle peak interferes with memory consolidation, while decoupling of the delta wave upstate and spindle peak augments it^7^.

When we examined the rate at which spindles nested within SO and delta wave upstates, we found a greater proportion of spindles coupled to delta waves than SOs, in line with the greater rate of delta waves (**Figure S3B–D**). We also found that WT TBI mice maintained temporally precise nesting of spindles to SOs even when the rate of both was increased after injury (**Figure S3C**). Of note, WT TBI mice had less contralateral spindle–delta wave nesting compared to the Sema3a mutant TBI mice, which themselves did not show any notable decrease in either SO–spindle nesting or delta–spindle nesting (**Figure S3C**).

### Sema3a preserves NREM spectral power after TBI

Finally, we measured the spectral power of various oscillatory frequencies (delta, theta, and sigma) present in both NREM and wake. Despite sustaining cortical tissue loss, WT TBI mice maintained intact ipsilateral spectral power in key NREM rhythms, including delta (0.3–5 Hz), theta (5–9 Hz), and sigma (9–15 Hz), relative to the contralateral hemisphere (**Figure 2A–C**). By contrast, Sema3a mutants exhibited reduced spectral power ratios in all 3 frequency bands after TBI (**Figure 2A–C**). These spectral effects were most pronounced in NREM; in wake, only a modest reduction in the sigma power ratio was observed in the Sema3a mutant TBI mice compared to WT TBI mice, with no significant change in delta or theta power ratios (**Figure S4A–C**).

**Figure 2.**
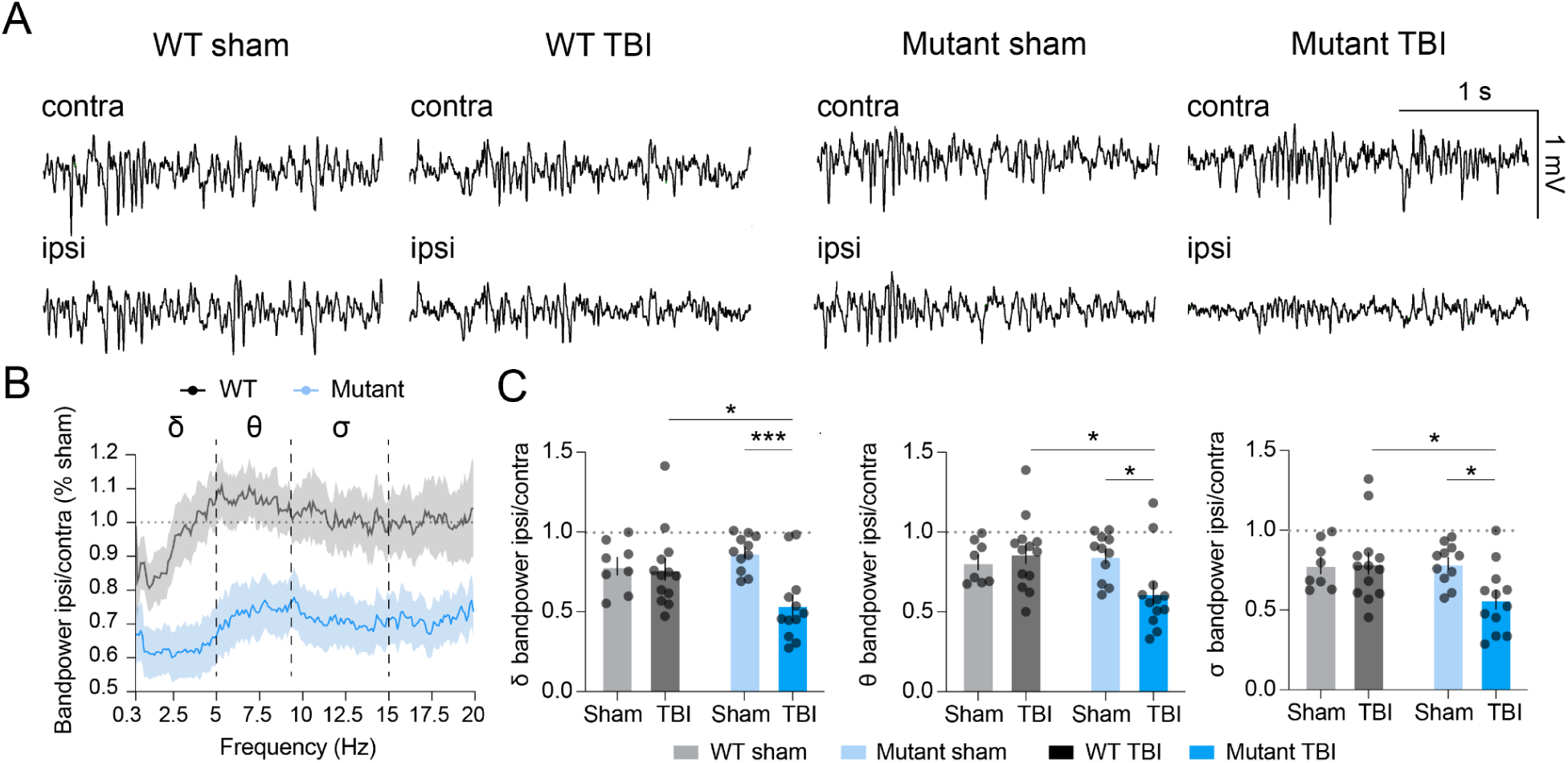
Sema3a preserves NREM spectral power after TBI. **(A)** NREM contralateral (contra) and ipsilateral (ipsi) traces from representative mice. **(B)** Ipsi/contra power in NREM normalized by respective shams. WT sham, N = 8; WT TBI, N = 14; Mutant sham, N = 11; Mutant TBI, N = 13. **(C)** Quantification of delta (δ), theta (θ), and sigma (σ) power in B. N’s as in B. Two-way ANOVA with Holm–Sidak multiple comparisons correction. \**p* < 0.05, \*\*\**p* < 0.001. Error bars show mean ± SEM across animals.

### Sema3a is required for sleep-related spatial memory consolidation and spontaneous behavioral reorganization after TBI

Because only WT TBI mice exhibited pro-memory shifts in NREM oscillations and the preservation of NREM spectral power, we next asked whether Sema3a signaling is required for memory consolidation 1.5 months after injury (Figure 3A).

**Figure 3.**
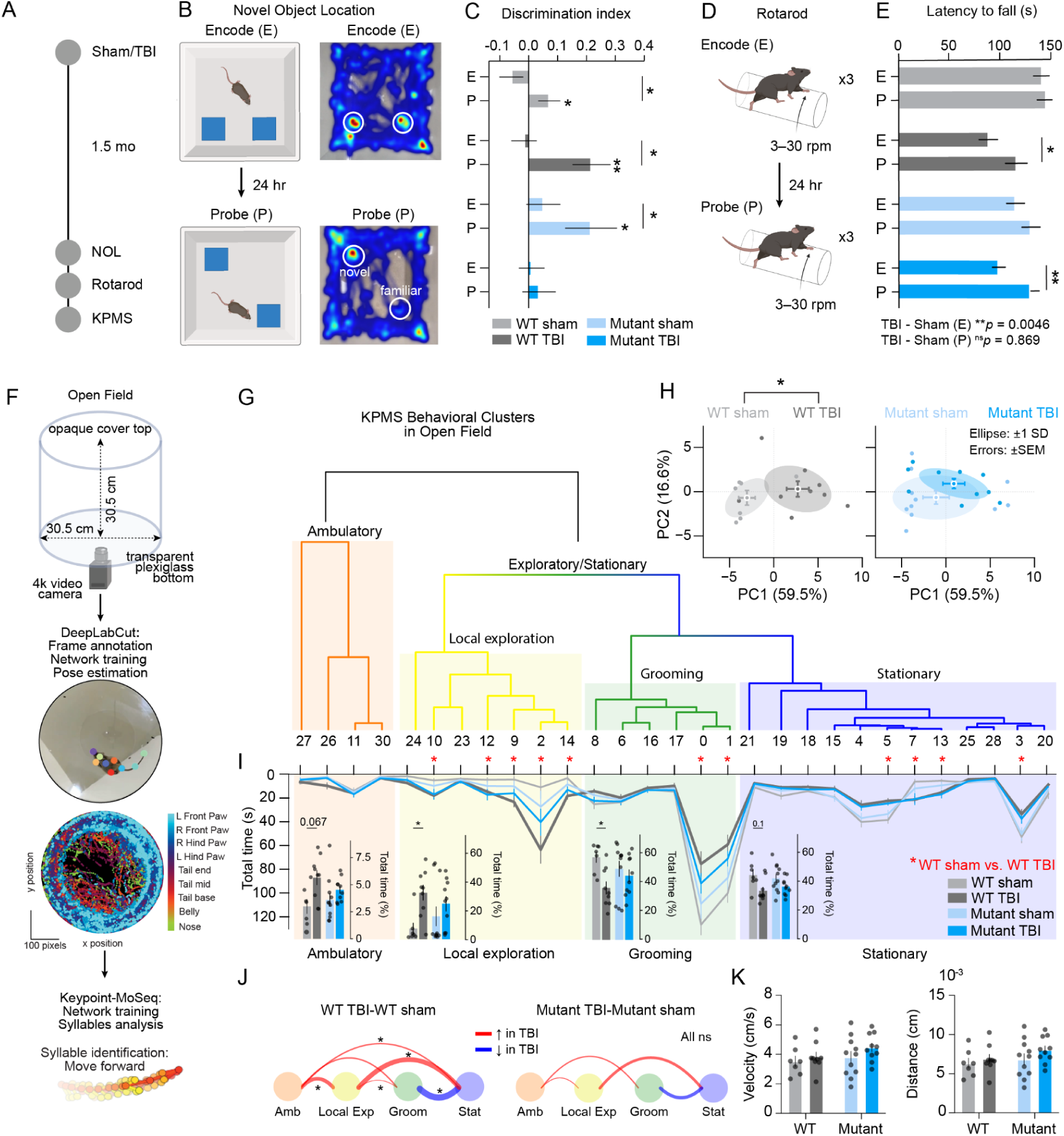
Sema3a is required for sleep-related spatial memory consolidation and spontaneous behavioral reorganization after TBI. **(A)** Experimental timeline of behavioral tests performed 1.5 months post-injury: NOL = Novel Object Location, Roto = accelerated rotarod, KPMS = Keypoint-MoSeq. **(B)** Schematic of NOL setup showing encoding (E) and probe (P) trials with representative heatmaps of mouse nosepoint position. Image created with Biorender.com. **(C)** Discrimination index over E and P trials. Paired two-tailed t-tests compare E vs P trials (asterisks connecting bars). One-sample two-tailed t-tests compare discrimination index to 0 (asterisks on individual bars). WT sham, N = 14; WT TBI, N = 17; Sema3a mutant sham, N = 18; Sema3a mutant TBI, N = 17. **(D)** Schematic of the accelerated rotarod task. Image created with Biorender.com. **(E)** Average latency to fall in E and P trials. Two-way ANOVA with Holm–Sidak multiple comparisons correction. Paired two-tailed t-tests to compare E and P. N’s as in C. **(F)** Open field apparatus and KPMS analysis pipeline. **(G)** Hierarchical clustering dendrogram of behavioral syllables organized into four higher-order clusters: ambulatory, local exploration, grooming, and stationary. **(H)** Principal component analysis of behavioral syllable frequencies. PC1 (59.5% variance explained) captures TBI-induced behavioral reorganization. PC1 and PC2 score distribution with individual mice shown as points (top) and group centroids ± SEM (bottom). Two-way permutation ANOVA (5,000 permutations) with Holm-Sidak multiple comparisons correction. WT sham, N = 7; WT TBI, N = 9; Mutant sham, N = 11; Mutant TBI, N = 10. **(I)** (Line plot) Total time (s) spent in each behavioral syllable, arranged by dendrogram cluster order. Red asterisks indicate a significant TBI effect in WT mice. Two-way permutation ANOVA (5,000 permutations) with Benjamini-Hochberg FDR procedure multiple comparisons correction (α = 0.05). Post-hoc comparisons used permutation t-tests with Holm-Sidak multiple comparisons correction. (Bar graphs) Percent of total 30-minute session spent in each behavioral cluster. Two-way permutation ANOVA (5,000 permutations) with Holm-Sidak multiple comparisons correction. N’s as in H. \**p* < 0.05, \*\**p* < 0.01. Error bars show mean ± SEM across animals. **(J)** Transition plots between behavioral clusters are shown as the difference between WT sham and WT TBI, and between Mutant sham and Mutant TBI. Red lines indicate an increased number of transitions in TBI relative to sham, whereas blue lines indicate decreased transitions. Line thickness reflects the magnitude of the difference between sham and TBI groups. Mann-Whitney permutation tests (5,000 permutations) with Benjamini-Hochberg FDR procedure multiple comparisons correction (α = 0.05). N’s as in H. **(K)** Average velocity and total distance traveled in the open field arena over the session. N’s as in H. \**p* < 0.05, \*\**p* < 0.01. Error bars show mean ± SEM across animals.

We evaluated spatial memory consolidation using a novel object location (NOL) task with a 24-hour delay, a paradigm highly dependent on sleep and functionally linked to SO and spindle activity^11,29–31^. The task involves two phases: an encoding phase, where mice explore two identical objects, followed 24 hours later by a probe phase, during which one object is moved to a new location (**Figure 3B**). WT sham, WT TBI, and Sema3a mutant sham mice all showed a preference for the novel object location in the probe phase, indicating intact spatial memory consolidation (**Figure 3C**). By contrast, Sema3a mutant TBI mice exhibited no novelty preference (**Figure 3C**).

Next, we tested motor memory consolidation using an accelerated rotarod task with a 24 hour delay (**Figure 3D**). During the encoding phase, latency to fall was recorded as mice ran on a constantly accelerating rod (**Figure 3D**). The mice were retested 24 hours later during the probe phase. Notably, injury impaired performance in the encoding phase but did not impede putative offline gains (performance improvement after sleep) in motor memory in either genotype (**Figure 3E**).

To further characterize the role of Sema3a in the behavioral consequences of TBI, we used Keypoint-MoSeq (KPMS), an unsupervised machine-learning approach that identifies discrete behavioral “syllables” during spontaneous open field exploration **(Figure 3F**, **Table 2)**. Hierarchical clustering of the 31 identified syllables revealed four primary behavioral clusters: ambulatory, local exploration, grooming, and stationary behaviors (**Figure 3G**).

**Table 2.**
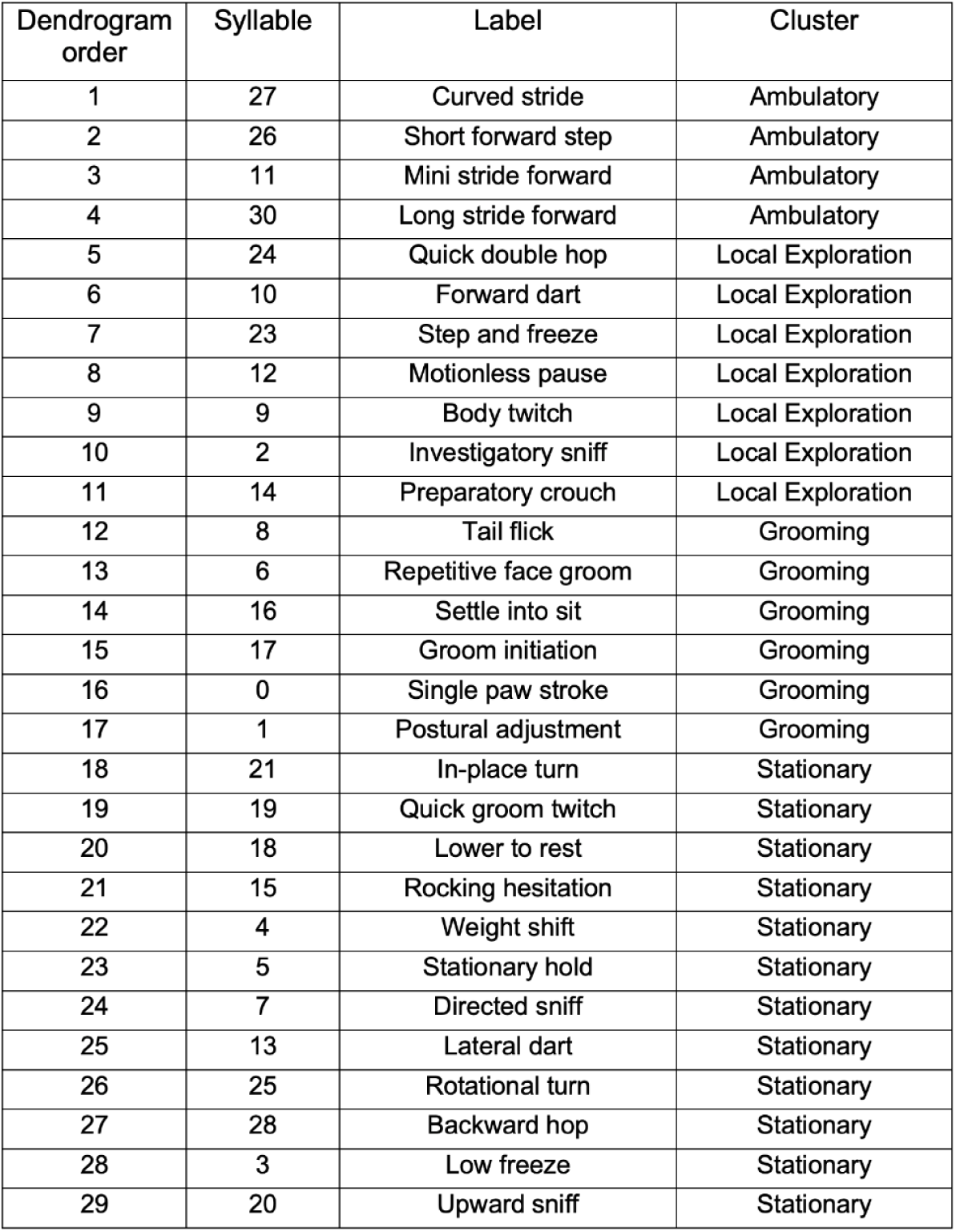
Syllable characterization for Keypoint-MoSeq.

To assess whether TBI induced global changes in motor behavior architecture, we performed principal component analysis (PCA) of syllable frequencies. PC1 captured 59.5% of the variance and clearly separated TBI from sham animals, although the effect was significant only in WT mice **(Figure 3H)**. Moreover, only WT mice exhibited a significant TBI-induced shift in spontaneous motor behavioral architecture **(Figure 3I)**. At the individual syllable level, TBI significantly altered the time spent in 11 of 31 syllables in WT mice, including increased time in local exploration behaviors and decreased time in grooming behaviors **(Figure 3I)**. By contrast, Sema3a mutant mice did not show significant TBI effects in any syllable **(Figure 3I)**. WT TBI mice also transitioned more frequently between behavioral clusters than WT sham mice, whereas Sema3a mutant TBI mice showed transition patterns similar to their sham controls **(Figure 3J)**. Neither genotype exhibited any overall change in average velocity or distance traveled (**Figure 3K**).

Collectively, these findings indicate that Sema3a selectively promotes spatial memory consolidation and reorganization of spontaneous behavior architecture after TBI, without affecting motor memory consolidation. This dissociation is consistent with the preferential dependence of spatial memory on hippocampal–thalamocortical circuits and the sleep oscillations they support, compared to the cortico-striatal circuits that predominantly underlie motor learning^30,32–34^.

### Sema3a limits cortical lesion expansion but not thalamic neuronal loss after TBI

Because TBI can lead to progressive neurodegeneration even years after the initial insult, we asked whether the lack of Sema3a signaling might affect **cortical lesion size at both acute and** chronic timepoints **(Figure 4A)**^35^. While cortical lesion volumes were comparable between WT TBI and Sema3a mutant **TBI mice** at 1–3 days after injury, Sema3a mutant TBI mice exhibited larger cortical lesions at 3 months compared to both 1**–**3 day WT TBI and 1**–**3 day Sema3a mutant TBI mice (**Figure 4C**). By contrast, 3 months after injury, WT TBI mice had lesion volumes equivalent to those at the 1–3 day timepoint (**Figure 4C**). Thus, Sema3a may protect against progressive cortical **lesion expansion.**

**Figure 4.**
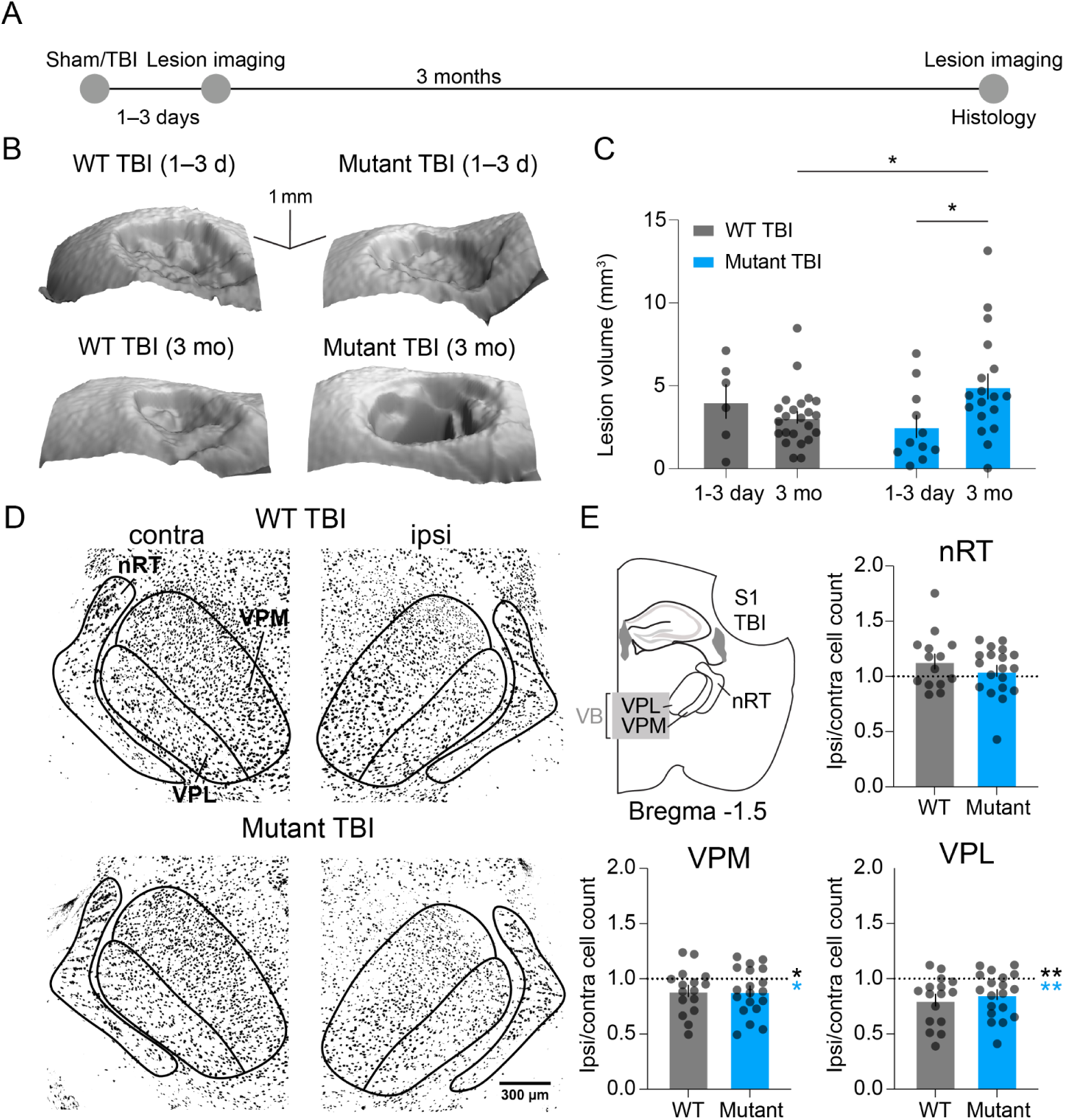
Sema3a limits cortical lesion expansion but not thalamic neuronal loss after TBI. **(A)** Experimental timeline **(B)** Representative images of the cortical lesion of a WT TBI mouse and a Sema3a mutant TBI mouse at 1–3 days post-injury and 3 months post-injury. Scale bar, 1 mm x 1 mm x 1 mm, **(C)** Quantification of the lesion volume at 1–3 days post-injury and 3 months post-injury. 1–3 days: WT TBI N = 6; Sema3a mutant TBI mice N = 11. 3 months: WT TBI N = 23; Sema3a mutant TBI N = 18. Two-way ANOVA with Holm–Sidak multiple comparisons correction. **(D)** Representative images of NeuN in the contralateral (contra) and ipsilateral (ipsi) thalami in a WT TBI and Sema3a mutant TBI animal. Scale bar, 300 µm. **(E)** Schematic of brain section with relevant regions demarcated and ipsi/contra cell count in the thalamic reticular nucleus, ventroposterolateral (VPL), and ventroposteromedial (VPM) nuclei of WT TBI and Mutant TBI mice. WT TBI n = 14 nRT, N = 8; Sema3a mutant TBI n = 19, N = 10; WT TBI VPL n = 16, N = 8; Sema3a mutant TBI VPL n = 19, N = 10; WT TBI VPM n = 16, N = 8; Sema3a mutant TBI VPM n = 19, N = 10. Asterisks next to the ratio = 1 dotted line indicate ipsi vs. contra comparisons (Wilcoxon matched-pairs signed-rank test); asterisks next to bars indicate WT vs. Sema3a mutant comparisons (Mann–Whitney test). \**p* < 0.05, \*\**p* < 0.01. Error bars show mean ± SEM across sections.

The thalamus—including the *nucleus reticularis thalami* (nRT), also known as the thalamic reticular nucleus, and the ventrobasal thalamus (VB)—is highly vulnerable to secondary injury and, together with the cortex, orchestrates oscillations critical for sleep and sleep-dependent cognition, including sleep spindles and SOs ^14,18,22,36–41^. Therefore, we also measured thalamic neuronal loss at 3 months post-injury and found that, while the nRT exhibited no change in NeuN+ cell counts between the contralateral and ipsilateral hemispheres, both the ventroposterolateral (VPL) and the ventroposteromedial (VPM) nuclei, which together compose the VB, had a reduced number of NeuN+ cells independent of genotype (**Figure 4D and 4E**).

## DISCUSSION

### Sema3a drives adaptive NREM architectures and preserves memory consolidation after TBI

Here, we show that Sema3a is a critical endogenous factor required for adaptive changes in sleep architecture and behavior after TBI.

NREM sleep is enriched in oscillations such as SOs, delta waves, and sleep spindles, that have been implicated in memory consolidation and participate in injury recovery^7,8,12,16,42–45^. Namely, SOs and sleep spindles are thought to enhance memory, while delta waves, and especially isolated delta waves, are thought to impede it^7,12^. At the chronic timepoint, WT TBI mice but not Sema3a mutant mice had more bilateral SOs and contralateral sleep spindles alongside fewer bilateral total delta waves and a lower proportion of bilateral isolated delta waves (**Figure 1F–I, Figure S2A–D**). WT TBI mice also preserved ipsilateral power in key NREM rhythms like delta, theta, and sigma, in contrast to Sema3a mutant TBI mice (**Figure 2A-–C**). The selective preservation of spatial memory in WT TBI mice (**Figure 3C and 3D**) paralleled the adaptive changes in their NREM architecture. Together, these data position Sema3a as a critical endogenous mechanism supporting network–level resilience to injury.

### How might injury-induced changes in NREM architecture promote resilience?

Low-frequency activity is commonly observed after brain injury, so much so that it is often considered a clinical biomarker^15,46^. While such activity may include a mixture of SOs and delta waves, there is evidence that augmented SO activity, in particular, reflects an adaptive recovery process. Prior work in stroke models shows that enhanced SOs and SO–spindle coupling can support functional recovery, presumably by creating a permissive environment for adaptive neuroplasticity^12,16^. Consistent with these findings, increased slow wave activity in chronic TBI patients (>1 year post–injury) was associated with improved learning on an overnight memory consolidation task, particularly when injury was severe^47^. That SO and delta wave rate—presumed to play contrary roles in memory^7^—were modulated in opposite directions after TBI in our model suggests a coordinated tuning of sleep-related rhythms to promote recovery.

Importantly, SOs and delta waves retained spindle coupling in WT TBI mice (**Figure S3B–D**). This suggests that, despite changes in their rate after injury, they maintain the coordination typical of healthy sleep architecture.

Furthermore, WT TBI mice exhibited a contralateral increase in sleep spindle activity concurrent with increased slow oscillations, suggesting co-modulation consistent with shared thalamocortical generators (**Figure 1F and Figure S2A,D**). Because sleep spindles are known to correlate with enhanced learning and memory^33,48^, our findings suggest that, much like the post–injury changes in SOs and delta waves, increased spindle activity may reflect a neural recalibration that supports cognitive recovery.

### Disentangling the role of Sema3a in memory and spontaneous behavior

Our study revealed a selective vulnerability of spatial memory to loss of Sema3a function. In particular, we observed that Sema3a mutant TBI mice lacked novel object location preference in the NOL task (**Figure 3C**), whereas motor memory consolidation appeared to be intact in both genotypes (**Figure 3D**).

Both the kind of spatial memory tested in NOL and the motor memory test on the accelerated rotarod have been shown to benefit from SOs and spindles that occur during sleep^11,29–31,49–51^. Therefore, why might one task appear to be more Sema3a-dependent than the other? One possible explanation is that the NOL task depends more on the hippocampus than the accelerated rotarod task, which relies heavily on the striatum and motor cortex^30,32–34^. Indeed, hippocampal memories appear to be particularly dependent on sleep, possibly due to sleep-dependent hippocampal replay and hippocampally generated sharp-wave ripples, which are themselves coordinated with SOs and sleep spindles^9,33,52,53^. Finally, sleep may play a larger role in multi-step motor tasks that demand more extensive training^7,54,55^. Utilizing more complex motor tasks may reveal a previously unrecognized role for Sema3a in sleep-dependent motor memory consolidation.

Intriguingly, WT TBI mice showed a marked change in spontaneous behavior in the open field arena that was absent in Sema3a mutant TBI mice (**Figure 3F—J**). These behavioral changes occurred even though average velocity and total distance traveled in the open field were not significantly altered by genotype or injury, indicating a reorganization of behavioral structure in WT TBI rather than a global change in locomotor output (**Figure 3K**). The fact that oscillatory changes, memory preservation, and behavioral reorganization all follow the same pattern, presenting in WT TBI mice and absent in Sema3a mutant TBI mice, suggests they arise from a unified Sema3a-dependent response to injury.

### Remaining questions

Notably, Sema3a mutant sham mice differed statistically from their WT counterparts across most electrophysiological measures. Given Sema3a’s well-established roles in axon guidance and circuit assembly^56^, constitutive loss of Sema3a function could yield subtle baseline phenotypes. For example, Chipman and colleagues reported no baseline electrophysiological differences in the mouse hippocampus but noted a minor reduction in spontaneous miniature end-plate potential (mEPP) amplitudes at the neuromuscular junction (NMJ)^4^. Outcomes may therefore depend on regional differences in Sema3a function and expression^57^. Our findings also raise the question of whether Sema3a mutants experience a “ceiling” effect that limits injury-induced plasticity. Nevertheless, the within-genotype changes towards a resilient state were unique to WT mice, and they largely correlated with lesion volume only in WT mice.

## CONCLUSION

In summary, our findings identify Sema3a as an endogenous factor that preserves sleep architecture and spatial memory consolidation after brain injury. By coordinating network-wide adaptations, Sema3a may serve as a molecular substrate for the resilience of sleep and cognition after TBI.

## METHODS

All protocols were approved by the Institutional Animal Care and Use Committee at the University of California, San Francisco, and Gladstone Institutes. Male and female homozygous Sema3a^K108N^ mutant mice (*C3;B6-Sema3a^m808Ddg^/J* (*Mus Musculus*), The Jackson Laboratory,RRID:IMSR_JAX:014646) in which Sema3a cannot signal through its receptor neuropilin-1, were used for experiments at between 10 and 30 weeks of age, alongside WT littermate controls^4,58^. Genotyping was performed via Transnetyx. Comparable numbers of male and female mice were included in each experiment. Precautions were taken to minimize stress and the number of animals used. Mice were group-housed with same-sex littermates and fed and watered ad libitum. One mutant sham animal had a skull perforation and was excluded from all analyses.

### Controlled cortical impact (TBI)

We anesthetized mice with 4–5% isoflurane and placed them in a stereotaxic frame. During the surgery, mice were maintained at 1.5–3% isoflurane. We performed a 3 mm craniotomy over the right somatosensory cortex (S1) centered at −1 mm posterior to Bregma and +3 mm lateral to the midline. TBI was performed with a CCI device (Impact One Stereotaxic Impactor for CCI, Leica Microsystems) equipped with a metal piston using the following parameters: 3 mm tip diameter, 20° angle, depth 0.8 mm from the dura, velocity 3 m/s, and 100 ms dwell time^36^. Sham mice received identical anesthesia and post–surgery care but no craniotomy, except sham animals that underwent ECoG recording, which were craniotomized. Immediately following surgery, mice were treated with a topical antibiotic and 5% lidocaine (McKesson Corporation) at the incision site, sterile saline intraperitoneally (8.0 mg/mL) (Fisher Scientific, Thermo Fisher Scientific Inc.), and extended-release analgesic buprenorphine (3.25 mg/kg) (Ethiqa XR, Fidelis Animal Health) delivered subcutaneously in the neck.

### Chromogenic DAB immunohistochemistry and microscopy

We anesthetized mice with a lethal dose of Fatal-Plus and perfused with 4% PFA in Ca^2+^/Mg^2+^free 1X PBS. Brains were transferred from 4% PFA to 30% sucrose 24 hours after perfusion and allowed to sink to the bottom (1–2 days), after which serial coronal sections 30 µm thick were cut on a Leica SM2000R microtome. Floating sections containing the relevant thalamic areas (∼Bregma 0.9 mm to 1.9 mm) were collected in ice-cold cryoprotectant (50% ethylene glycol / 1X PBS / 20% glycerol) and stored at –20°C. All incubation steps were performed with gentle agitation.

Sections were briefly rinsed 3 times with 0.1 % Tween 20 in PBS (PBS-Tw), then incubated for 20 minutes in fresh 1% H2O2 in 30% Ethanol/PBS. After another 3 PBS-Tw rinses, sections were blocked in 1X PBS-Tw containing 20% normal goat serum and Mouse-on-Mouse IgG Blocking Reagent (Vector Laboratories, MKB-2213-1) (2 drops/2.5 ml block solution) for 1 hour at RT. After blocking, the sections were incubated overnight at 4 °C with anti-NeuN (1:500, mouse, Sigma-Aldrich, MAB377, AB_2298772) diluted in 1X PBS-Tw with 3% normal goat serum.

After 3 5-minute 1X PBS-Tw washes, sections were incubated with MACH 2 Mouse HRP-Polymer (Biocare Medical, MHRP520L) for 1 hour. After another 3 5 minute 1X PBS-Tw washes, sections were developed with the DAB/Ni Peroxidase Substrate Kit (Novus Biologicals, SK-4100-NB) for approximately 1 minute and 30 seconds. After a brief rinse in 1X PBS, sections were mounted. After sections dried completely, they were dehydrated in the following sequence: 70% ethanol (1 x 1 minute), 95% ethanol (2 x 1 minute), 100% ethanol (2 x 2 minutes), Xylene (2 x 2 minutes). Finally, sections were coverslipped with Epredia Shandon-Mount (Fisher Scientific, 1900333) and stored long-term at 4 °C.

For each brain section, a 10X tile scan was collected for an overview of the sample on a Keyence BZ-X810. NeuN puncta were counted in ImageJ/FIJI using a custom macro. Images were thresholded, converted to 8-bit grayscale, and binarized, then subjected to watershed segmentation to separate adjacent objects. Puncta were defined using a minimum size threshold of 50 pixels with circularity constrained between 0.2 and 1.0. Only particles fully contained within the region of interest (ROI) were included, and final counts were normalized to the ROI area.

### Surgical implantation of electrocorticography (EcoG) and electromyography (EMG) devices

ECoG signal was acquired using a custom-made Millmax-based device^14^. Cortical screws were implanted bilaterally in somatosensory cortex S1 (+0.5 mm anterior from Bregma, ± 3 mm lateral) and symmetrically in the contralateral hemisphere. A reference screw was implanted above the cerebellum (−0.5 mm posterior to Lambda, 0.5 mm lateral to the midline). The ipsilateral S1 cortical screws were implanted 1 mm anterior to the most anterior edge of the craniotomy. A contralateral stability screw was also placed approximately symmetrical to the center of the craniotomy to help the device remain fixed to the skull. A stripped wire was used to record electromyographic (EMG) activity and was placed in the trapezius muscle of the neck^59,60^. Devices were fixed to the skull using dental cement (Henry Schein). Immediately following surgery, mice were treated with lidocaine, saline, and buprenorphine as previously described (see “Controlled cortical impact (TBI)”).

### In vivo electrocorticography (ECoG) data acquisition

To control for circadian rhythms, animals were housed using a regular light/dark cycle, and recordings were performed between 7:00 AM and 7:00 PM. Mice were allowed to recover for one week prior to the recording. ECoG signals were recorded using RZ5 via Synapse software (Tucker Davis Technologies) and sampled at 1017 Hz. Animals were continuously monitored during recordings using a video camera that was synchronized to the signal acquisition via the RZ5^59,60^. Recordings were 4 hours in duration. A 0.3–50 Hz bandpass single-pole digital filter was applied during acquisition. All analyses were conducted on the filtered data.

### Sleep staging

ECoG data from the contralateral hemisphere were utilized for sleep staging. First, non-physiological periods of activity exceeding 1 mV in either the positive or negative direction were excluded. Sleep staging was performed automatically using a custom Python function that identifies putative NREM epochs based on overlapping periods of elevated delta power and reduced muscle activity; epochs were defined by the intersection of EMG RMS and ECoG delta power thresholds.

Features relevant for sleep staging were calculated in 5-s segments. Z-scored EMG RMS was calculated within each window and compared to the median and standard deviation of the RMS distribution of the entire noise–free signal. Windows with z-scored RMS values below the threshold (mean of RMS) were retained as candidate NREM periods. In parallel, z-scored delta power (0.3–5 Hz) was calculated per window. Windows with delta power exceeding a threshold (mean +0.5 SD) were classified as putative NREM. Only windows that satisfied both criteria were retained, and overlapping segments were used to define the final NREM epochs. Any epoch not classified as NREM was classified as wake. Because features like SOs and sleep spindles are most prominent in NREM, REM epochs were excluded. Gaps shorter than 30 s between consecutive NREM epochs were concatenated, and valid epochs needed to be at least 10 seconds in length. Final epochs were manually inspected. In a few cases, excessive noise precluded accurate sleep segmentation; depending on the level of noise, NREM was either manually defined or the entire recording was excluded. Animals that slept for less than 5 minutes over the course of the recording were excluded from further analysis due to insufficient sample size.

### Power analysis

To quantify spectral power in ECoG recordings, we used Welch’s method to compute the power spectral density (PSD) with a default Hann window over either concatenated NREM or wake epochs. For all analyses, the PSD was computed over the frequency range 0.3–20 Hz to capture the frequencies of greatest relevance. Absolute power for each frequency band (e.g., delta, theta) was calculated by integrating the PSD over the desired frequency range using Simpson’s rule. The frequency bands were defined as follows, exclusive of the end frequency: delta (0.3–5 Hz), theta (5–9 Hz), sigma (9–15 Hz)^37,61–63^.

### Sleep spindles

Sleep spindles were detected using a Morlet wavelet–based approach. A complex Morlet wavelet was generated at a center frequency of 15 Hz with 6 cycles to match the expected frequency and temporal characteristics of cortical spindles. Lower central frequencies were tested, but 15 Hz was optimal for detection. The wavelet was then convolved with the NREM ECoG signal to extract the analytic amplitude and phase. The instantaneous power was calculated by squaring the amplitude of the resulting signal.

Putative spindles were initially identified as contiguous periods where the power exceeded five standard deviations above the mean baseline power. To refine spindle boundaries, we extended the start and end of each event until the power dropped below a more permissive threshold of 3 standard deviations above the mean ±250 milliseconds on either side to capture the entire spindle envelope, including jitter around the threshold. Spindles were further filtered based on their duration (minimum: 0.8 s; maximum: 4 s) and minimum inter-spindle interval (0.5 s). Events separated by gaps shorter than this interval were concatenated into single spindle events so long as the spindles did not exceed the maximum length.

### Slow oscillations/delta + nesting

Slow oscillations (SOs) and delta waves were identified in NREM as previously described in Kim et al.^7^. Because sleep spindles tend to nest in wave upstates, upstates (representative of synchronous neuronal depolarization) were verified by the probability of spindle nesting. If a spindle’s peak amplitude in the 9–15 Hz filtered band (second-order bandpass Butterworth filter) occurred within 1 second of an SO upstate or a delta wave upstate, it was considered nested to the respective wave. To define isolated vs coupled delta waves, we calculated the temporal delay between a delta wave and its preceding SO (Δδwave–SO) and used the median of each distribution at the contra electrode to define the distinction between a coupled (Δδwave–SO < median) or isolated (Δδwave–SO > median). WT TBI and Sema3a mutant TBI used the medians of their respective shams (as described in the figure legend).

### Novel object location (NOL)

NOL was conducted between 7 AM and 7 PM in an opaque white acrylic 60 cm x 60 x 30 cm arena. At the start and end of each day, the arena was thoroughly bleached. In between animals, the arena was cleaned with 70% ethanol to minimize olfactory cues. All mice were tested in the same order and were placed in the center of the arena to commence each trial. Before the encoding and probe trials, mice were allowed to explore the empty arena freely for 10 minutes. 24 hours later, at the same time of day, mice were returned to the arena in the presence of two plastic blocks affixed to the arena floor 13 cm from the most proximal sides and were allowed to explore freely for 10 min. Object location in the encoding trial was counterbalanced. Mice were once again replaced in the arena 24 hours later, at the same time of day, with one block moved to a novel location. Mice were allowed to explore freely for 10 min. The mouse’s location was monitored via a video camera (SONY HDR-CX675) and analyzed using tracking software (Noldus, SCR_004074). A 5-cm-radius circular zone surrounded each object, and nosepoint entry into a zone for ≥1 second was counted as object exploration. The discrimination index was calculated as follows:

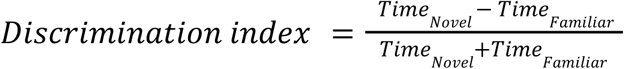

where exploration time was the sum of interactions with both objects during the encoding or probe trial. One WT TBI animal interacted with only one object during encoding and was excluded from DI analysis due to insufficient exposure, but was retained in total exploration time analyses.

### Accelerated rotarod

Accelerated rotarod testing was conducted between 7 AM and 7 PM, 24 hours after the mice underwent the NOL test. At the start and end of each day, the rods and landing platform were thoroughly bleached. Between animals, the rods and landing platform were cleaned with 70% ethanol to minimize olfactory cues. All mice were tested in all behavioral tests in the same order. Mice were loaded on the rotarod (Med Associates Inc. ENV-574R) while the rod (7 cm diameter, 11.5 cm width, 43 cm fall height). rotated at a constant rate of 3 rotations per minute (rpm). The rotarod was then programmed to accelerate from 3 to 30 rpm over 300 seconds. Each trial terminated when the mouse fell off the rod or at 300 seconds. Mice were replaced on the rotarod, and each run was repeated for a total of 3 trials for the encoding trial. 24 hours later, the experiment was repeated for the probe trial. Average latency to fall was calculated across the 3 trials in each session.

### Keypoint-MoSeq

#### Pose Estimation

Mice were placed in a circular open field (12-inch/30.5-cm diameter) with a clear bottom. They were video-recorded from below using a 4K video camera (Logitech) at 48 frames per second (fps) for 30 minutes. Twenty frames per recording were labeled for the following key points: nose, all four paws, belly, and the base, middle, and end of the tail. These labeled images were used to train a DeepLabCut (DLC; RRID:SCR_021391) network for 1 million iterations^64,65^. Pose trajectories were derived from egocentric coordinates obtained through DLC pose estimation, with the nose as the anterior reference point and the tail base as the posterior reference point. The tail midpoint and tail tip were excluded from downstream analysis due to high tracking variability. For egocentric alignment, the anterior reference point was determined using the nose, while the posterior reference point was identified as the base of the tail. A principal component analysis (PCA) of these egocentrically-aligned key point time series was extracted from DLC and used to create a model in Keypoint-MoSeq (RRID:SCR_025032)^66^.

#### Syllable Extraction

Following the standard workflow in the KPMS repository (https://github.com/dattalab/keypoint-moseq/), we fit an autoregressive hidden Markov model (AR-HMM) to the pose trajectory data, setting the hyperparameter kappa to 10,000,000. The AR-HMM was fit over 50 iterations, followed by a full KPMS model trained for 500 iterations. KPMS output of frame-by-frame syllable usage was used to generate frequency and duration distributions for each mouse. The model identified 31 behavioral syllables from frame-by-frame pose sequences. Syllables 22 and 29 occurred at very low frequencies and durations and lacked representative .gif and .mp4 outputs; these syllables also did not cluster coherently within the similarity dendrogram and were therefore excluded.

#### Syllable Labeling and Classification

Each syllable was assigned a descriptive label by an observer who reviewed the corresponding .gif animations, .mp4 video clips, and their associated pose trajectory time series. A similarity dendrogram of the 29 syllables was used to identify four overarching behavioral clusters (ambulatory, local exploration, grooming, and stationary), named based on the predominant behavioral syllables in each cluster.

### Lesion volume

Images of the lesion were obtained using the Keyence VK–X1000 3D laser-scanning microscope (SCR_027341) and analyzed using the accompanying MultiFileAnalyzer software^67^. The reference plane for volume was set to a flat cross-section outside the injury. If the brain had both concave and convex injuries, the absolute volumes of both were summed to obtain the total lesion volume. The area around the lesion was manually drawn and selected for analysis. The height threshold was set to the highest point for concave and the lowest point for convex to account for the entire volume of the lesion. Any brains whose lesion sites were damaged during perfusion were excluded. Moisture artifacts were removed from representative images using Adobe Photoshop for clarity, but only the raw data were analyzed.

### Statistical analyses

All numerical values are given as means, and error bars are the standard error of the mean (SEM) unless stated otherwise. The threshold for statistical significance was set to α = 0.05. \**p* < 0.05,\*\**p* < 0.01,\*\*\**p* < 0.001,\*\*\*\**p* < 0.0001. n refers to slices, sections, or cells where applicable, while N refers to the number of mice. Normality was assessed using the D’Agostino & Pearson, Anderson–Darling, Shapiro–Wilk, and Kolmogorov–Smirnov tests. Parametric and nonparametric tests were chosen as appropriate and are reported in the figure legends. Post-hoc comparisons were only made within genotype and injury families unless otherwise stated (i.e., comparisons were not made between WT sham and Mutant TBI, nor between Mutant sham and WT TBI). Data analysis was performed with Clampfit 10.7 (RRID: SCR_011323), GraphPad Prism 10 (SCR_002798), ImageJ/FIJI (SCR_003070), and Spike2 (SCR_000903) software.

## KEY RESOURCES TABLE

| REAGENT or RESOURCE | SOURCE | IDENTIFIER |
| --- | --- | --- |
| Antibodies |  |  |
| Mouse anti-NeuN, clone A60 (1:500 IHC) | Sigma-Aldrich | RRID:AB_2298772 |
| Critical commercial assays |  |  |
| DAB/Ni Peroxidase Substrate Kit | Novus Biologicals | Cat#SK-4100-NB |
| Experimental models: Organisms/strains |  |  |
| *C3;B6-Sema3a<sup>m808Ddg/J</sup> (Mus Musculus)* | The Jackson Laboratory | RRID:IMSR_JAX:014646 |
| Software and algorithms |  |  |
| ImageJ/FIJI | NIH | RRID: SCR_003070 |
| Spike2 | Cambridge Electronic Design | RRID: SCR_000903 |
| GraphPad Prism 10 | GraphPad | RRID: SCR_002798 |
| Other |  |  |
| MACH 2 Mouse HRP-Polymer | Biocare Medical | Cat#MHRP520L |
| Mouse-on-Mouse IgG Blocking Reagent | Vector Laboratories | Cat#MKB-2213-1 |

## RESOURCE AVAILABILITY

### Lead contact

Further information and requests for resources and reagents should be directed to and will be fulfilled by the lead contact, Jeanne Paz.

### Materials availability

This study did not generate unique reagents.

### Data and code availability

All original data is available upon request from the lead contact, Jeanne Paz.

## ACKNOWLEDGMENTS

We would like to thank Dr. Grae Davis and Dr. Pete Chipman for invaluable scientific discussions as well as the gifting of the Sema3a ^K108N^ mutant mice. We would also like to thank Drew Willoughby and Jeremy Ford for analysis assistance, the Biomolecular Nanotechnology Center (BNC) at UC Berkeley for access to the Keyence VK–X1000 3D laser-scanning microscope, and the Gladstone Histology Core for access to the Keyence BZ-X810.

## AUTHOR CONTRIBUTIONS

Conceptualization, D.N. and J.T.P;

Formal analysis, D.N., Y.V., S.P.;

Methodology and visualization, D.N., Y.V., S.P.;

Software, D.N. and Y.V.;

Writing – original draft, D.N.;

Writing – review and editing, D.N., Y.V., J.T.P.;

Funding acquisition, J.T.P.

---

**Supplementary Figure 1.**
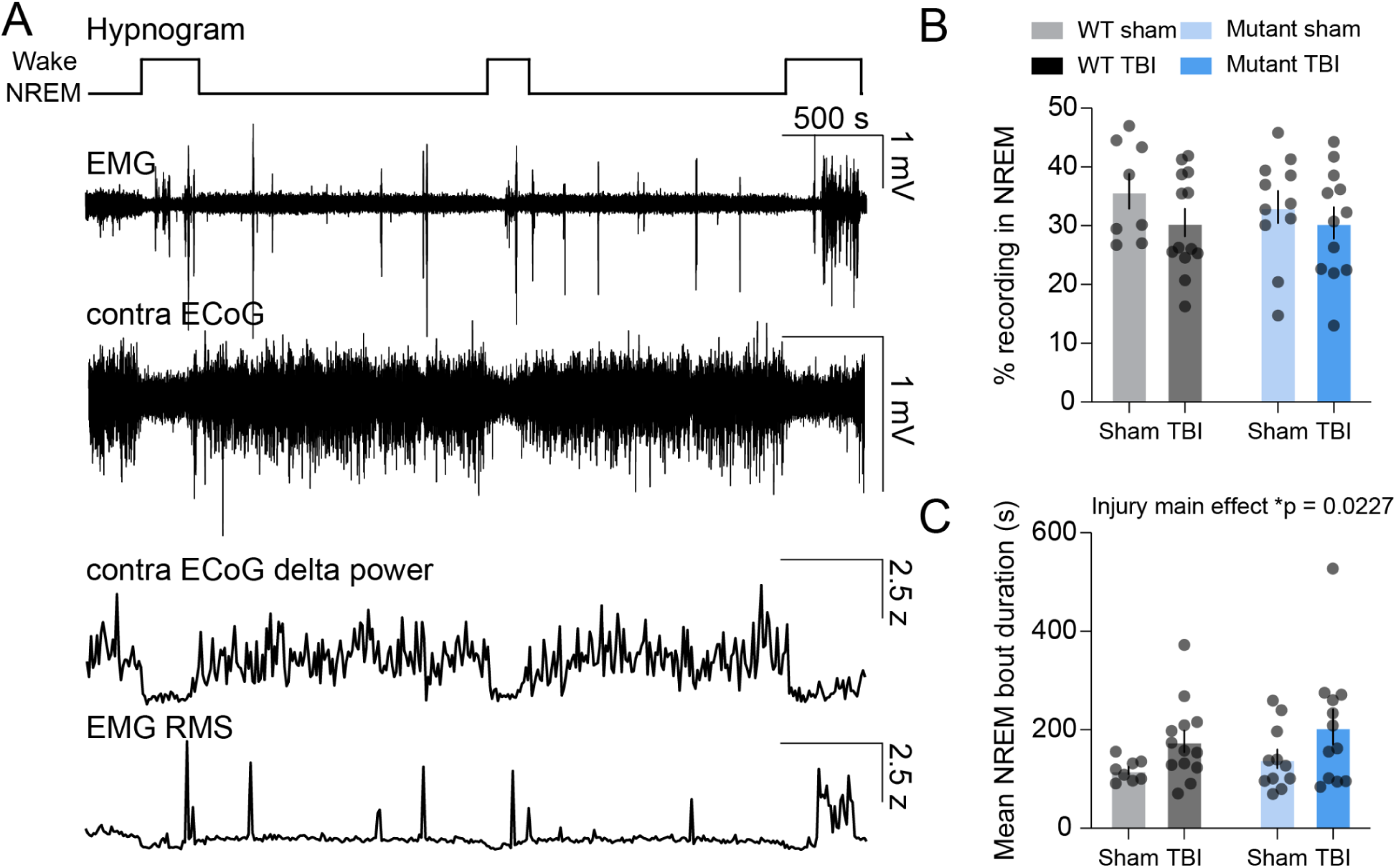
TBI increases NREM bout duration independent of Sema3a after TBI. **(A)** Representative hypnogram, EMG, contralateral (contra) ECoG (0.3–20 Hz), contra ECoG delta power (0.3–5 Hz), and contra EMG root mean square (RMS). Pink lines indicate thresholds above and below which delta power and EMG RMS, respectively, would qualify for NREM, provided other criteria were met (see Methods). Z in scale refers to the z-score. **(B-C)** Percent of total recording in NREM (B) and mean NREM bout duration (C). WT sham, N = 8; WT TBI, N = 13; Sema3a mutant sham, N = 11; Sema3a mutant TBI, N = 12. Two-way ANOVA with Holm–Sidak multiple comparisons correction. \**p* < 0.05. Error bars show mean ± SEM across animals.

**Supplementary Figure 2.**
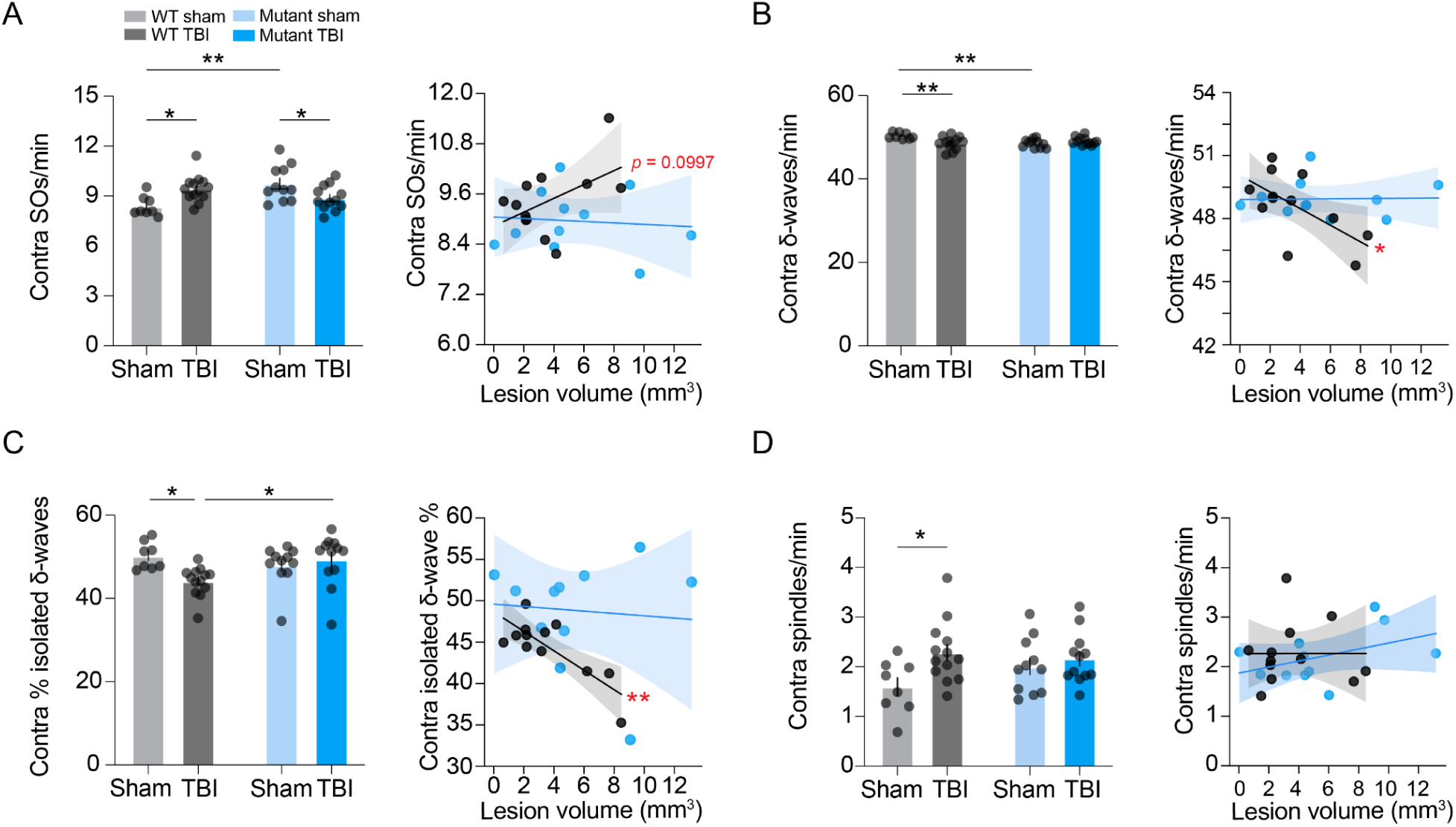
Sema3a is required for contralateral adaptive tuning of NREM architecture after TBI. **(A–D)** Slow oscillation (SO) and delta wave (δ-wave) rate in NREM on the contralateral (contra) hemisphere with lesion volume correlations. WT sham, N = 8; WT TBI, N = 14; Sema3a mutant sham, N = 11; Sema3a mutant TBI, N = 13. Two-way ANOVA with Holm–Sidak multiple comparisons correction. Simple linear regression for correlations. Asterisks next to each regression line indicate a significant *p*-value. See Table 1 for linear regression statistics.

**Supplementary Figure 3.**
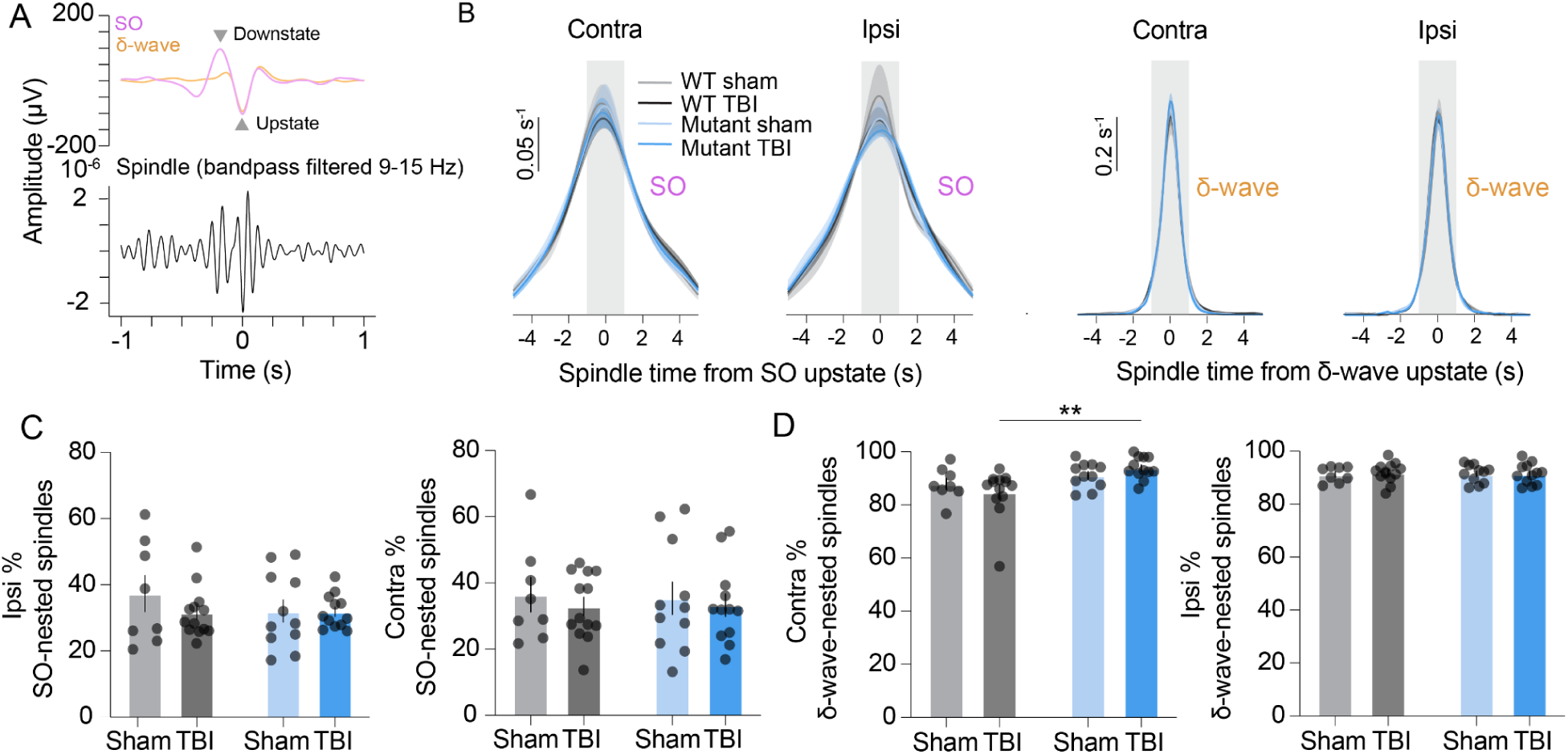
Sleep spindle-SO and delta wave nesting is preserved after TBI. **(A)** Average slow oscillations (SOs) and delta waves (δ-waves) from a representative mouse with the average 9–15 Hz filtered ECoG. **(B)** Probability density functions of the delay between the SO or δ-wave upstate and the nearest spindle peak. The gray box indicates the time window (± 1 s from wave upstate) in which a spindle is considered nested. WT sham, N = 8; WT TBI, N = 13; Sema3a mutant sham, N = 11; Sema3a mutant TBI, N = 12. **(C)** Percent of spindles nested to SOs. N’s as in B. Two-way ANOVA with Holm–Sidak multiple comparisons correction. **(D)** Percent of spindles nested to δ-waves. N’s and statistics as in C. \**p* < 0.05, \*\**p* < 0.01. Error bars show mean ± SEM across animals.

**Supplementary Figure 4.**
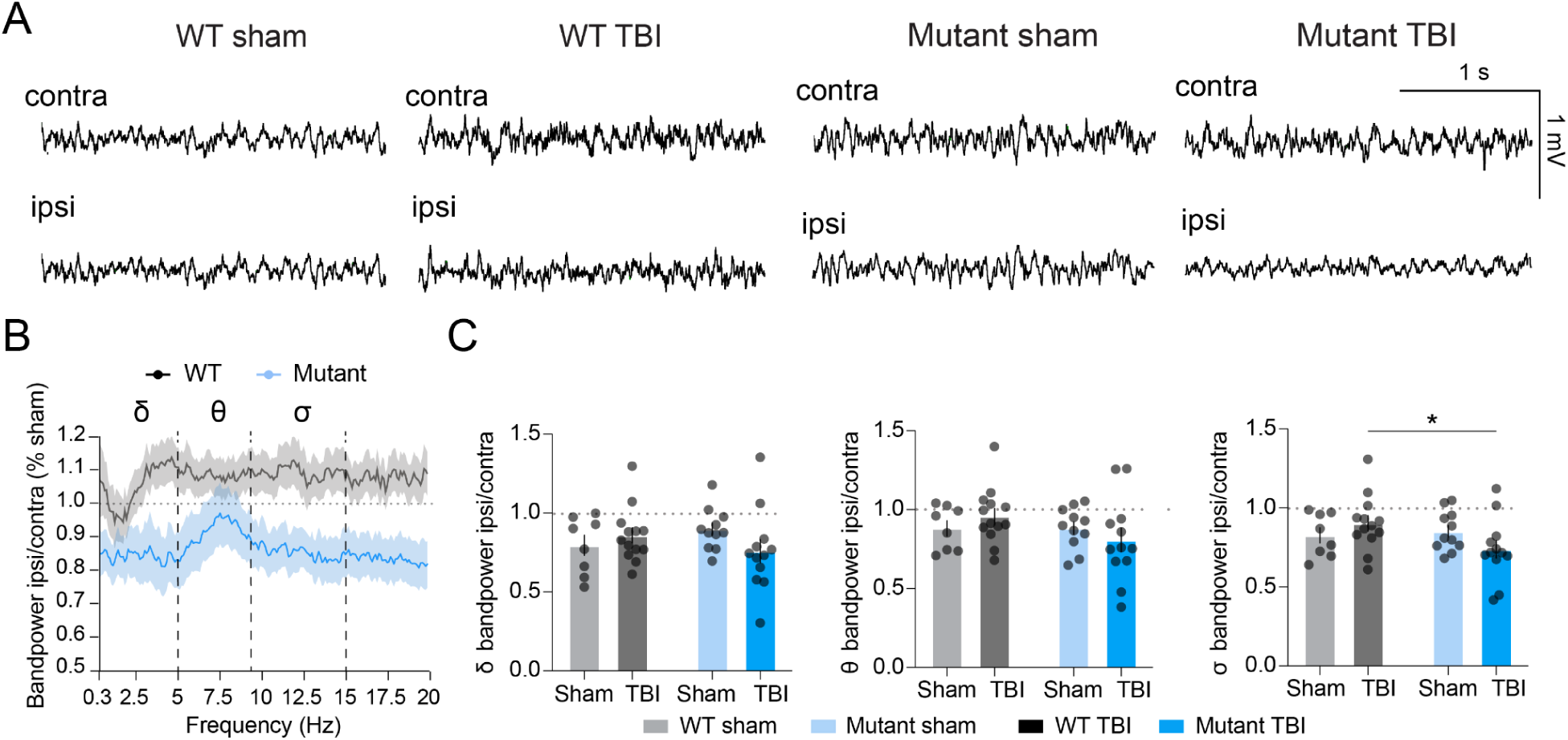
Sema3a moderately preserves wake bandpower after TBI. **(A)** Wake contra and ipsi traces from representative mice. **(B)** Ipsi/contra power in wake normalized by respective shams. WT sham, N = 8; WT TBI, N = 13; Mutant sham, N = 11; Mutant TBI, N = 12. **(C)** Quantification of delta (δ), theta (θ), and sigma (σ) power from B. N’s as in B. Two-way ANOVA with Holm–Sidak multiple comparisons correction.

